# Developing a Standard Definition for Sequences of Concern

**DOI:** 10.64898/2026.03.14.711820

**Authors:** Tessa Alexanian, Jacob Beal, Craig Bartling, Jens Berlips, Peter A. Carr, Adam Clore, Helena Cozzarini, James Diggans, Yorgo El Moubayed, Kevin Esvelt, Kevin Flyangolts, Leonard Foner, Patrick A Fullerton, Bryan T Gemler, Caitlin AD Jagla, Rassin Lababidi, Tom Mitchell, Steven T Murphy, Michael T Parker, Nicholas Roehner, Andre Rusch, Kemper Talley, Troy Timmerman, Nicole E Wheeler

## Abstract

Readily available nucleic acid synthesis is both critical for the bioeconomy and an increasingly pressing security concern due to the potential for accidental or deliberate misuse. While biosecurity experts broadly agree that nucleic acid providers should screen orders for potential “sequences of concern,” there has previously been no agreed standard for how to define and recognize such sequences.

To address this gap, we first organized a test set of 1.1 million sequences from pathogens and toxins on the Australia Group Common Control Lists and their non-controlled relatives, along with model organisms and synthetic constructs. An initial categorization of sequences as to whether or not they were sequence of concern was produced by comparing the results of four biosecurity screening systems for each of these sequences, finding that these systems already agreed on the categorization of more than 80% of sequences.

We then refined these results through a science-based stakeholder review process to define a rubric for determining whether a sequence should be flagged as a potential sequence of concern, then applied this rubric to improve the categorization of test sets. The result is a rubric that identifies sequences of concern with respect to human pandemic-potential viruses, key classes of low-risk genes, and controlled toxins. Applying this rubric to the test set collection has reduced the number of test sequences with disputed categorization by 44.3% for controlled viruses and 10.7% across the test set as a whole. Together, these results provide a concrete “sequence of concern” definition that can be used as a foundation for development of biosecurity screening standards and policy.

## 1 INTRODUCTION

Readily available nucleic acid synthesis is both a critical supply chain input for the bioeconomy and an increasingly pressing security concern, given the potential for accidental or deliberate misuse of genes for dangerous pathogens or toxins (Hoffmann et al., 2023; Diggans and Leproust, 2019). There is broad consensus amongst biosecurity stakeholders that one of the key defenses against such misuse is screening of nucleic acid orders for sequences of concern (World Health Organization, 2022; of Sciences Engineering et al., 2018; International Gene Synthesis Consortium, 2024; International Organization for Standardization, 2024), and this recommendation has begun to be incorporated into biosecurity policy, including in both the United States (The White House Office of Science and Technology Policy (OSTP), April 2024; The White House, 2025) and the UK (UK Department for Science, Innovation, and Technology, 2024). For screening requirements to be effective, however, it is necessary to ground them in shared standards for evaluating the efficacy of screening methods, which in turn requires at least some degree of agreement on which sequences should be considered “sequences of concern” (Wheeler et al., 2024a; Laird et al., 2025; Millett et al., 2023).

Accordingly, the Sequence Biosecurity Risk Consortium (SBRC) was founded to address this need by bringing together synthesis providers, screening tool developers, policymakers, and scientific experts to create and maintain a science-based definition for sequences of concern (Beal and Alexanian, 2025). The SBRC grew directly out of the International Gene Synthesis Consortium’s (IGSC) Test Set Working Group, which developed the multi-tool comparison methodology used here to develop draft categorizations (Wheeler et al., 2024a). The IGSC also maintains the Regulated Pathogen Database (International Gene Synthesis Consortium, 2024) that serves as the taxonomic foundation for the test sets. This evolution from an industry-led working group to a broader multi-stakeholder consortium reflects the IGSC’s longstanding role in establishing biosecurity screening as an industry norm, now formalized into a process capable of supporting enforceable standards.

Here we report on the first key results from this effort: development of a large-scale sequence test set and a complementary rubric for categorization of sequences. We first generated a large-scale test set comprising approximately 1.1 million sequences from pathogens and toxins on the Australia Group Common Control Lists and their non-controlled taxonomic relatives, along with model organisms and synthetic constructs. We next determined the initial categories of test set sequences through comparison of tool results using the methodology established in our prior work (Wheeler et al., 2024a), finding an overall agreement on the categorization of more than 80% of test sequences. These results were then refined through through a science-based stakeholder review process adapted from the community standards development processes used by the IETF (Resnick, 2014), Python (Warsaw et al., 2000), and SBOL (SBOL Community, 2025) communities.

The product of this work, the SBRC Screening Testing Collection version 1.0 reported herein, is a rubric that defines sequences of concern with respect to human pandemic-potential viruses, key classes of low-risk genes, and controlled toxins, along with the application of that rubric to reduce the number of test sequences with disputed categorization by 44.3% for controlled viruses and 10.7% across the test set as a whole. Together, these results provide a concrete “sequence of concern” definition that can be used as a foundation for development of biosecurity screening standards and policy.

In the remainder of this paper, we first detail the construction of the test set and initial categorization of sequences in Section 2, then describe the refinement process and its results in Section 3. Finally, Section 4 summarizes these results and discusses how they can be used to support enforceable biosecurity screening requirements, as well as discussing ongoing efforts in the SBRC and needs for complementary standards development.

## 2 TEST SET CONSTRUCTION AND INITIAL CATEGORIZATION

We began the process of defining sequences of concern with an empirical approach, previously prototyped in Wheeler et al. (2024a), in which de facto areas of agreement and dispute are discovered by running sequences through multiple biosecurity tools and comparing their results. While agreement does not necessarily indicate a correct assessment of risk, areas of dispute are likely to be more important to resolve in order to ensure that dangerous orders can be consistently identified. We thus first constructed a large-scale test set intended to enable comprehensive testing, then evaluated all test set sequences in four biosecurity screening tools, and finally analyzed the results of this initial categorization as a candidate empirical definition of sequences of concern.

### 2.1 Test Set Construction

We began by scaling up the methodology used in Wheeler et al. (2024a) to construct a collection of test sequences covering the full range of pathogens and toxins in the IGSC Regulated Pathogen Database (International Gene Synthesis Consortium, 2024). The IGSC RPD covers all regulated biological agents listed on the Australia Group Common Control Lists, the US Export Administration Regulation Commerce Control List, the US Federal Select Agent Program, and the European Union list of dual-use items, comprising 39 taxonomic clusters of regulated pathogens, plus additional lists for regulated toxins.

For each taxonomic cluster, test sets were constructed for both regulated threats and closely related relatives. For example, the *Bacillus cereus group* cluster contains threat taxa *Bacillus anthracis* and *Bacillus cereus biovar anthracis* and non-regulated relatives such as *Bacillus cereus, Bacillus thuringiensis*, and *Bacillus toyonensis*. Test sets were constructed for nucleic acid sequences and reverse-translated protein sequences, each comprising up to 10,000 sequences in NCBI between 200 bp and 10,000 bp in length from the relevant taxa. In cases where more than 10,000 sequences were available, the number was reduced by taking a random subset of the available NCBI sequences.

Exceptions were made in the following cases: viroids do not express proteins, and thus have no protein test sequences; the *Peronosporaceae* cluster has no protein threat test sequences because NCBI currently contains no protein sequences besides a few highly conserved genes; and the toxin collections use additional filtering to remove non-toxin genes.

Finally, two additional clusters of low-risk sequences were added to address common sources of false positives: one containing 10,000 bp sequences taken from the genomes of a diverse collection of model organisms and one containing engineered nucleotide sequences drawn from the iGEM registry of standard biological parts (iGEM Foundation, Accessed March, 2026).

The result is a collection of approximately 1.1 million sequences divided into 42 clusters organized into four “kingdom-level” categories: 13 viral clusters, 17 bacterial clusters, 7 fungal clusters, and 5 other clusters—viroids, two toxin collections, model organisms, and engineered sequences.

### 2.2 Screening Tool Comparison

To make an initial determination about whether each sequence should be considered a sequence of concern, we applied the workflow from Wheeler et al. (2024a), as shown in Figure 1. Each sequence was independently screened by four biosecurity screening systems: Aclid (Aclid, Inc., Accessed March, 2026), Battelle UltraSEQ (Gemler et al., 2023), the IBBIS Common Mechanism (Wheeler et al., 2024b), and RTX BBN FAST-NA Scanner (Beal et al., 2021; Wyschogrod et al., 2022), each applying is own distinct and independently developed algorithmic approach to determining whether to flag a sequence as a potential sequence of concern.

**Figure 1.**
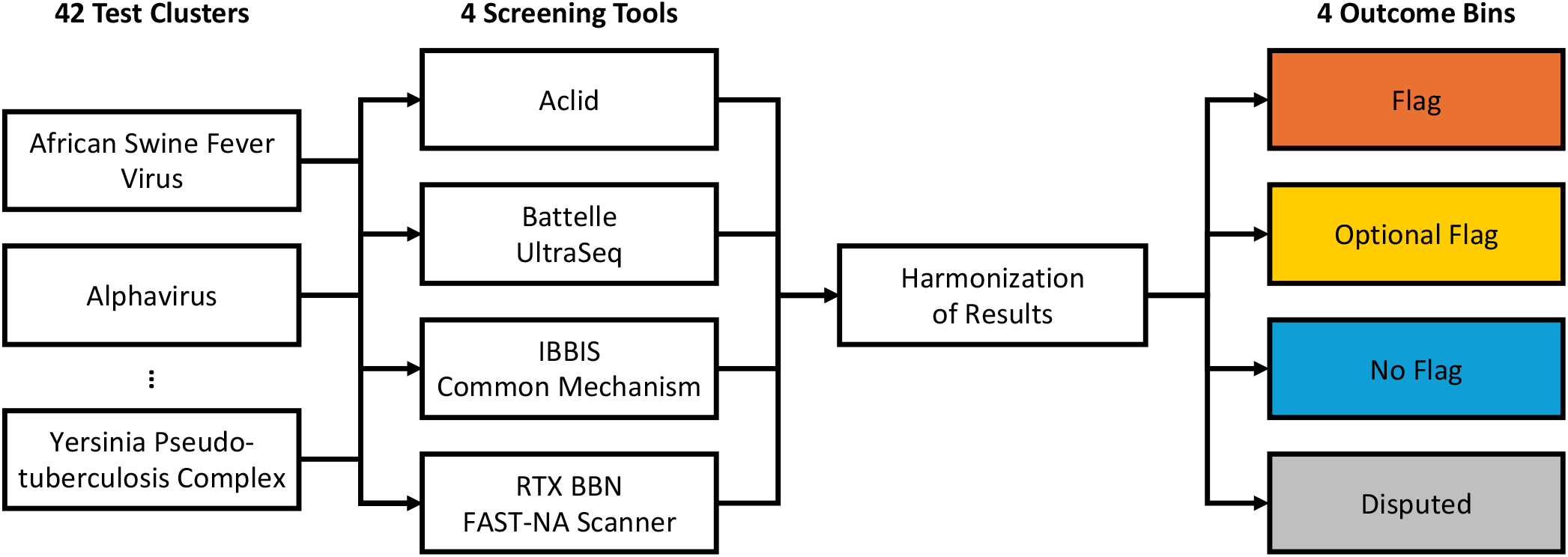
Initial categories for test sequences were determined by evaluating the sequences of each cluster independently with four biosecurity screening tools, then harmonizing results to categorize each sequence into one of four categories of consensus screening outcome. Figure adapted from Wheeler et al. (2024a).

The heterogenous reports produced by the four tools are then harmonized by mapping each tool’s result into one of three categories (“Flag”, “No Flag”, or “Optional Flag”) and combining the results following the decision logic shown in Figure 2 to produce one of four consensus screening outcomes:

**Figure 2.**
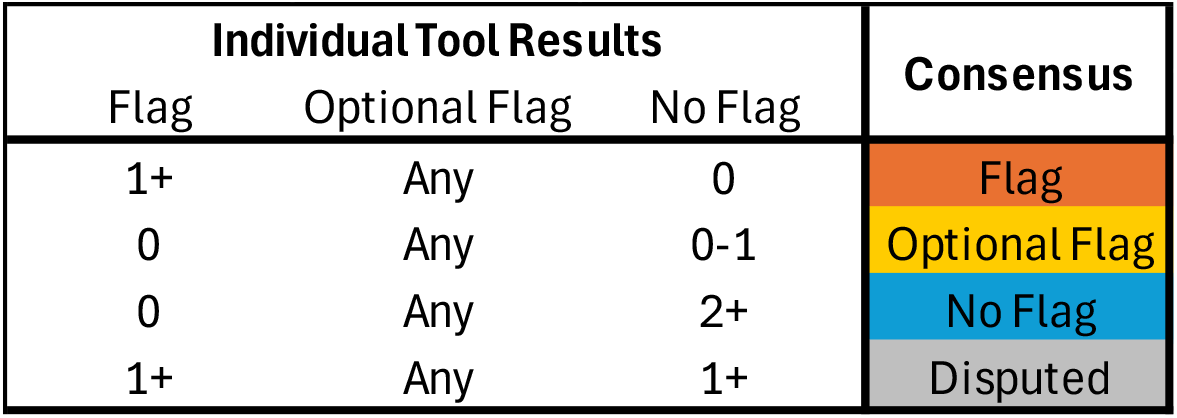
The consensus screening outcome across tools is determined by comparing the results of individual tools following the decision logic shown in this table.

- **Flag**: sequence is “guilty” of being a sequence of concern (i.e., agreed to be potentially dangerous, such that an order containing that sequence should be flagged for follow-up screening): flagged by at least one tool and not cleared by any tool.
- **No Flag**: sequence is “innocent” of being a sequence of concern (i.e., a low-risk sequence that should not be flagged): cleared by at least two tools and not flagged by any tool.
- **Optional Flag**: sequence is “not guilty” and “not innocent” (i.e., not enough evidence to decide on a category): cleared by zero or one tool and not flagged by any tool.
- **Disputed**: sequence has a “hung jury” of tools disagreeing on biosecurity risk: flagged by at least one tool and cleared by at least one tool.

Note that the “Disputed” category was previously termed “Undetermined” in Wheeler et al. (2024a), but has been relabelled to better reflect how sequences in the category are being interpreted.

### 2.3 Results of Initial Categorization

Figure 3 shows the results of the initial categorization produced by comparing the results of screening tools. Overall, the rate of agreement is high, with tools achieving either a “Flag” or “No Flag” consensus on 80.5% of all sequences. The rate of agreement is far higher for sequences from relatives than for threats, but even for threats there is agreement on the majority of sequences (52.8%). Importantly, the lower rate of agreement for threats is driven primarily by “Optional Flag” sequences for cellular threats (particularly fungi) rather than by “Disputed” sequences. This is consistent with the prior results in Wheeler et al. (2024a) and with the observation that many genes in these organisms are currently too poorly understood to make a meaningful assessment of potential risk, and thus also generally likely to be too poorly understood to be readily misused. Only 4.4% of sequences are “Disputed”: 8.5% of threats and 1.73% of relatives. Note that for relatives, some sequences were indeed flagged by some tools, but every such sequence was cleared by at least one other tool and thus ended up in the “Disputed” category.

**Figure 3.**
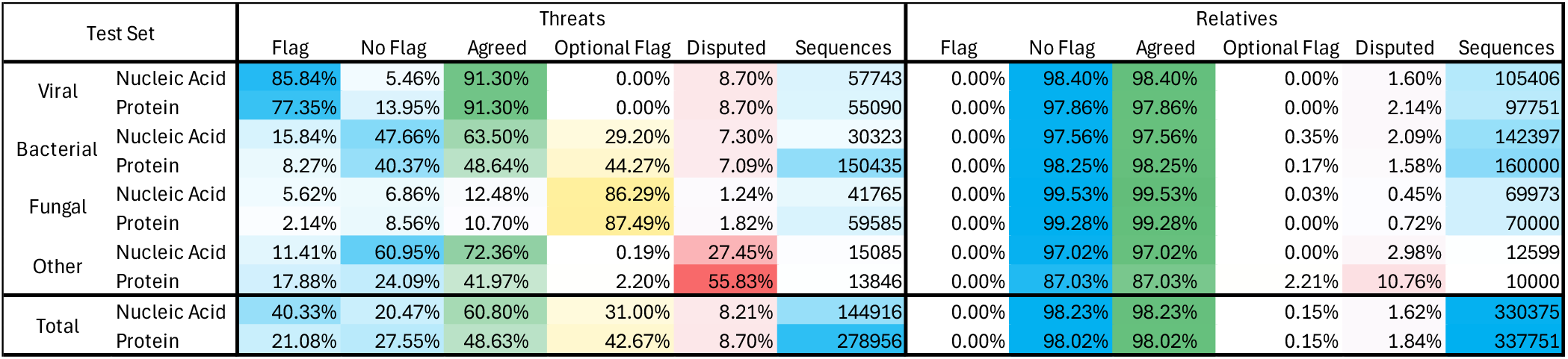
Results of initial categorization of test set sequences, with “Agreed” category summing “Flag” and “No Flag” results. Color intensity is used to highlight relative value.

At the level of kingdoms, results for threats are again consistent with the prior small-scale test in Wheeler et al. (2024a). Amongst the threat sequences, viruses have the highest level of agreement, bacterial sequences are intermediate, and fungal sequences are dominated by poorly understood “Optional Flag” genes. The highest levels of disputes are around certain classes of toxins for which there are disagreements between tools on how to define risk. For bacteria, disputes are driven primarily by the interpretation of genes, while for viruses disputes are driven more by the interpretation of taxonomy. Results for relatives are, as one would expect, consistently nearly all “No Flag”, with the exception of certain classes of toxin. Results for nucleic acid sequences and reverse-translated protein sequences are generally consistent with one another, with the largest differences being primarily a statistical effect of the difference in numbers of sequences between test sets.

All told, these results show that there already is a high degree of consistency between the implicit definitions of sequences of concern that are implemented by the biosecurity screening tools used to produce this initial categorization of sequences. At the same time, there is no inherent authority in the judgements of these particular tools. These results thus provide a starting point that needs to be complemented with a science-based approach to identifying and correcting errors and resolving disputes in the categorization of test set sequences.

## 3 STAKEHOLDER-DRIVEN REFINEMENT OF DEFINITIONS

The SBRC complements the “bottom up” empirical definition of sequence of concern in the test sets with a “top down” definition in the form of the Biosecurity Flag Rubric, a structured process for determining whether to flag a given nucleic acid sequence. The Biosecurity Flag Rubric and test sets are both updated by community processes for proposing and reviewing improvements. These processes have been applied by the SBRC community to refine the initial categorization of test sequences into an improved definition of sequences of concern.

### 3.1 Community Processes

The Biosecurity Flag Rubric is created and maintained through a community proposal process modeled on established practices used by other standards communities (Figure 4). In particular, the SBRC embraces decision-making based on the highly successful Internet Engineering Task Force (IETF) rough consensus process (Resnick, 2014), using proposal templates inspired by the Python community (Warsaw et al., 2000), and polls to affirm consensus modeled on those of the Synthetic Biology Open Language (SBOL) community (SBOL Community, 2025).

**Figure 4.**
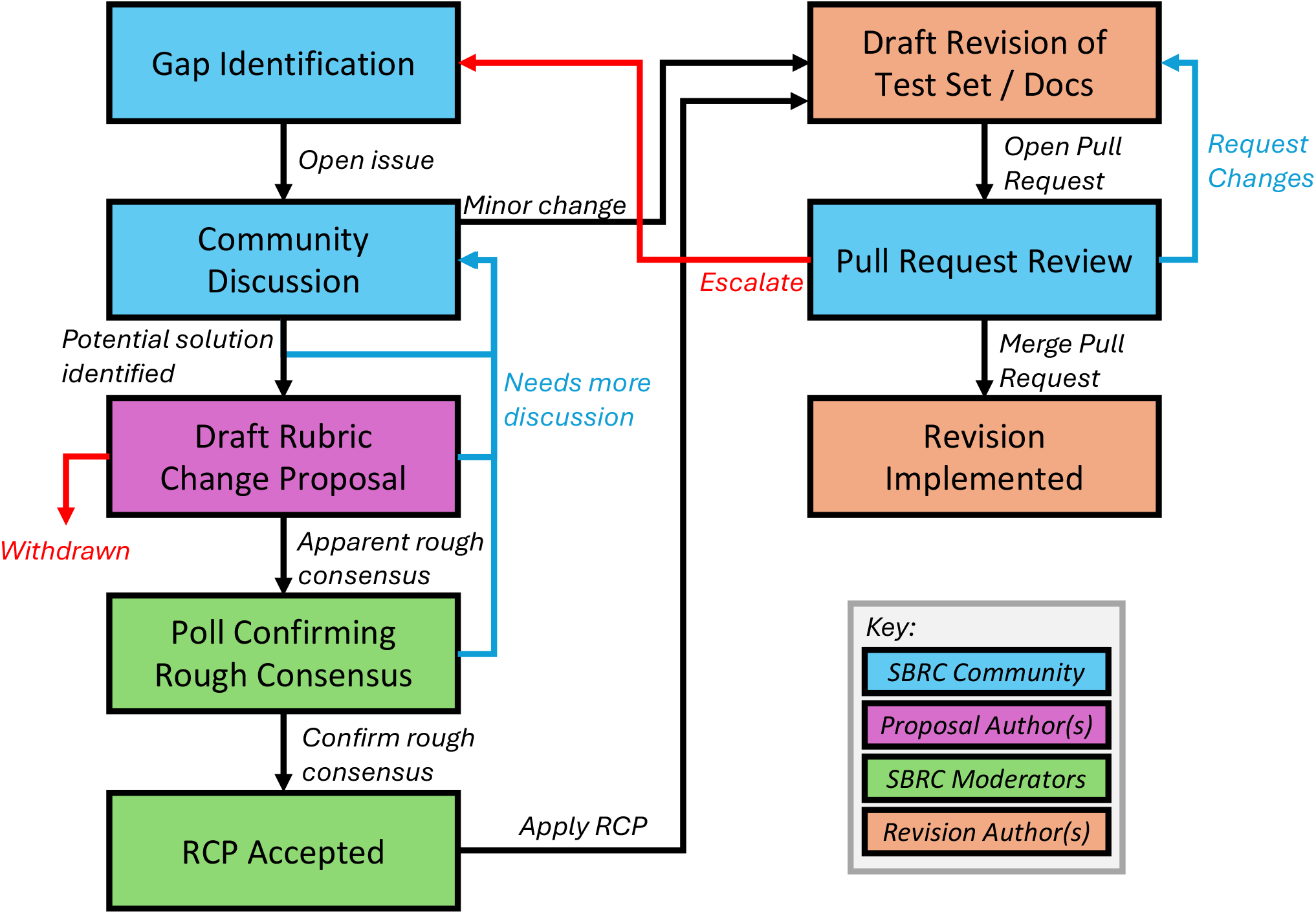
The SBRC revises the Biosecurity Flag Rubric and makes meta-level decisions through a process of IETF-style rough consensus (Resnick, 2014) on Rubric Change Proposals, then applies these decisions to revise test sets and documentation with the aid of pull request reviews.

Changes to the Biosecurity Flag Rubric and meta-level decision-making regarding community processes are handled via *Rubric Change Proposals* (RCPs) as follows:

- **Gap identification and community discussion:** Any member of the SBRC community who notices a gap or other concern needing to be addressed can open an issue for community discussion.
- **Community discussion:** The issue is discussed and potential solutions are put forth. If the necessary solution involves no real substantive changes (e.g., updating a biological name to track external resources or clarifying the wording used to describe a process), at this point it may be judged a “minor change” that does not need community review and can be fast-tracked directly to a draft revision and pull request (see below).
- **Draft Rubric Change Proposal:** Once a viable potential solution has been identified and seems to be gaining support amongst community members, it is made concrete by drafting a formal Rubric Change Proposal (RCP) providing the rationale for the proposed approach, the specific changes that are being proposed, and concrete examples of revisions that will result from adoption of the RCP. The community member who identified the gap often also authors the RCP, but there is no necessary relationship. Discussion continues and the RCP is revised as needed until it either appears to have achieved a state of rough consensus (i.e., no significant technical concerns remain that have not been addressed) or else is withdrawn.
- **Poll Confirming Rough Consensus:** Because much of the SBRC’s work is done asynchronously, it is important to ensure that all community members have the opportunity to review an RCP and raise concerns before it is accepted. For this reason, once the SBRC community moderators assess that an RCP appears to have achieved rough consensus, they run a formal poll with a one-week notification and comment period, followed by a one week period to collect responses. Because RCPs do not proceed to a poll until rough consensus is already believed to have been achieved, the typical outcome is unanimous responses in favor of acceptance. Any vote to the contrary is expected to be accompanied by an expression of specific technical concerns, which are either addressed on the spot if minor or else require the RCP to be returned to the community discussion stage.

Once a proposal has been accepted, it may be applied to revise test sets and documentation through a faster and more lightweight process based on standard agile software development practices. Any SBRC community member can author a revision to test sets and/or documentation. When the draft revision is ready, they submit it as a “pull request” to merge in the proposed revision (the term “pull request” is taken from software development, where it refers to a request to review and incorporate a proposed change). The draft is then reviewed by other community members, who either approve it to be merged, request changes to improve it, or escalate larger issues for consideration as a potential new RCP.

Taken all together, the result of this process is to ensure that changes to both the Biosecurity Flag Rubric and the test set are acceptable to a broad range of biosecurity stakeholders in the SBRC. The complete specification of this process is attached as Supporting Information: “RCP 001: Rubric Change Proposals.”

### 3.2 Results of Definition Refinement

Over the course of 11 months, the SBRC applied these processes to develop version 1.0 of the SBRC Screening Testing Collection, which was released privately in September 2025. During this period, the SBRC community put forward 23 RCPs, of which 17 were accepted and had changes incorporated into the release (implemented via dozens of pull requests), five were accepted and incorporated later, and one was withdrawn. Of the 17 accepted RCPs, four are process RCPs: one establishing the RCP process itself, one on community governance, and two establishing criteria for evaluating risk. The other 13 are Biosecurity Flag Rubric RCPs: one establishing the initial rubric, five addressing human pandemic-potential viruses, two identifying toxin genes, one identifying certain bacterial virulence factors, and four identifying classes of “No Flag” low-risk sequences.

The result of these activities is a Biosecurity Flag Rubric that defines sequences of concern with respect to human pandemic-potential viruses (8 of the 13 viral clusters), key classes of low-risk genes, and controlled toxins. Applying these definitions to the test sets allowed significant improvements in categorization of test set sequences, as shown in Figures 5 and 6. The primary area of improvement is in the viral test sets, where the five RCPs addressing human pandemic-potential viruses drove a 44.3%, reduction in the number of “Disputed” sequences for viral threats and an 18.1% reduction for their relatives. Combined with small reductions in “Disputed” sequence counts for other kingdoms from the other RCPs, this resulted in an overall reduction in “Disputed” sequences of 10.7% across the test set as a whole. There were also minor improvements in the number of “Optional Flag” sequences, but these were not a significant focus of effort (being a lower priority than disputes) and the overall reduction was only 0.34%.

**Figure 5.**
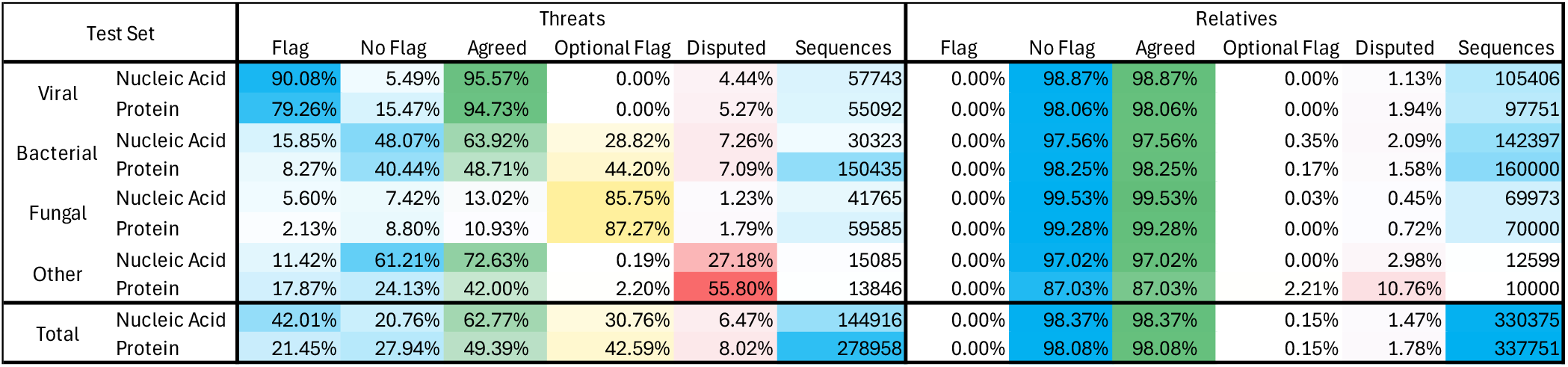
Categorization of test set sequences in SBRC Screening Testing Collection version 1.0, with “Agreed” category summing “Flag” and “No Flag” results. Color intensity is used to highlight relative value.

**Figure 6.**
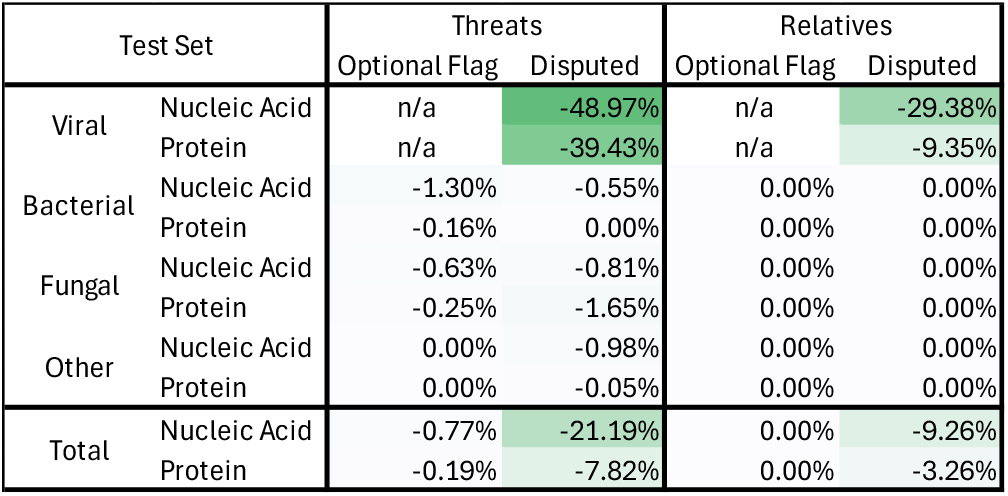
Reduction in number of “Optional Flag” and “Disputed” sequences in SBRC Screening Testing Collection version 1.0 relative to initial categorization. Color intensity is used to highlight relative value.

## 4 DISCUSSION

Together, the SBRC Biosecurity Flag Rubric and test sets provide a concrete “sequence of concern” definition that is known to be both acceptable to a broad range of biosecurity stakeholders and compatible with current industrial biosecurity screening best practices, coupled with a process for continuing to improve and maintain this definition over time. These results thus provide a solid foundation on which to build other components of effective biosecurity screening, including proficiency testing, certification, government regulation, and legal enforcement. Because the process grounds out in science-based risk assessment rather than regulation and is developed by a growing international community, it also offers a rendezvous point for international harmonization of standards and regulations. This is particularly timely given the current governmental efforts to develop nucleic acid screening regulations in the United States, United Kingdom, European Union, and other countries.

Beyond the work reported here, the SBRC has continued to develop the Biosecurity Flag Rubric and test sets. Ongoing efforts include expansion of the rubric to the full range of bacterial and fungal pathogens on the Australia Group lists, handling differences in regulation between different jurisdictions, extension to high-risk pathogens not currently regulated, function-based criteria for risk assessment, and resolution of disputed sequences. The SBRC is also developing systematic tests for screening evasion threats such as AI-assisted redesign of proteins (Wittmann et al., 2025), splitting-based obfuscation (Tayouri et al., 2025), and order fragmentation (Edison et al., 2026; Wittmann et al., 2026). The SBRC is also actively expanding its membership to incorporate additional subject-matter experts and stakeholders from a broader range of countries, with the aims of deepening the expertise available for science-based risk assessment (especially with respect to pathogens with limited geographic distribution) and increasing the opportunities for international harmonization.

It is also important to note that this work represents only one aspect of implementing effective enforceable biosecurity standards. Directly complementary to standards for defining sequences of concern is the need for customer screening standards that address the question of how to determine whether a customer should be given access to a sequence of concern (Mackelprang et al., 2025), including questions of customer pre-approval, verification, and the handling of “grey area” cases. Likewise, both governments and non-governmental organizations need to cooperate to utilize these standards to implement systems of certification, regulation, and legal consequences that can drive universal adoption of effective biosecurity screening practices by nucleic acid providers. Finally, the growing complexity of biotechnology and increasing capabilities of AI also point towards a need for biosecurity screening to be adopted more broadly across the whole of the bioeconomy at other points of potential intervention such as biological design tools, AI-assisted protein engineering software, and laboratory information management systems (LIMS), as well as Large Language Models (LLMs), agentic AI systems, and other general purpose tools with the potential to uplift biological capabilities.

## AUTHOR CONTRIBUTIONS

Conceptualization, Data Curation, Investigation, Methodology, Validation, Writing - review and editing: T.A., J.Bea., C.B., J.Ber., P.C., A.C., H.C., J.D., Y.E., K.E., K.F., L.F., P.F., B.G., G.G., C.J., R.L., T.M., S.M., M.P., N.R., A.R., K.T., T.T., N.W.

Formal Analysis, Software, Writing - original draft: J.Bea.

Project Administration, Resources: J.Bea., T.A., R.L.

## ACKNOWLEDGMENTS

Thank you for additional technical inputs to Jesse Bloom of the Fred Hutchinson Cancer Center and Gene Godbold of Signature Science. This document does not contain technology or technical data controlled under either U.S. International Traffic in Arms Regulation or U.S. Export Administration Regulations.

## DATA AVAILABILITY STATEMENT

The data used in this article are part of a curated research resource and are not publicly available. Access is granted to qualified organizations under licensing terms that mitigate potential information hazards and that support the maintenance and long-term sustainability of the data collection. Data access requests should be directed to the SBRC moderators. Further information about the data and associated licensing terms is available at https://sbrc.bio/.

## REFERENCES

Aclid, Inc. (Accessed March, 2026). Aclid. Online resource https://www.aclid.bio/

Beal, J. and Alexanian, T. (2025). Creating enforceable biosecurity standards for nucleic acid providers. Engineering Biology 9, e70003

Beal, J., Wyschogrod, D., Mitchell, T., Katz, S., Manthey, J., and Clore, A. (2021). Development and transition of fast-na screening technology. Tech. Rep. BBN Tech Report 8622

Diggans, J. and Leproust, E. (2019). Next steps for access to safe, secure dna synthesis. Frontiers in bioengineering and biotechnology 7, 86

Edison, R., Toner, S., and Esvelt, K. M. (2026). Assembling unregulated dna segments bypasses synthesis screening: regulate fragments as select agents. Nature Communications

Gemler, B. T., Mukherjee, C., Howland, C., Fullerton, P. A., Spurbeck, R. R., Catlin, L. A., et al. (2023). Ultraseq, a universal bioinformatic platform for information-based clinical metagenomics and beyond. Microbiology Spectrum 11, e04160–22

Hoffmann, S. A., Diggans, J., Densmore, D., Dai, J., Knight, T., Leproust, E., et al. (2023). Safety by design: Biosafety and biosecurity in the age of synthetic genomics. Iscience 26

iGEM Foundation (Accessed March, 2026). igem registry of standard biological parts. Online resource https://parts.igem.org

International Gene Synthesis Consortium (2024). Harmonized screening protocol v3.0. Online resource

International Organization for Standardization (2024). ISO 20688-2. Biotechnology—Nucleic Acid Synthesis, Part 2: Requirements for the Production and Quality Control of Synthesized Gene Fragments, Genes, and Genomes. Tech. rep., ISO

Laird, T. S., Flyangolts, K., Bartling, C., Gemler, B. T., Beal, J., Mitchell, T., et al. (2025). Inter-tool analysis of a nist dataset for assessing baseline nucleic acid sequence screening. Applied Biosafety, 15356760251401228

Mackelprang, R., Rivera, S., Klonowski, J., Smith, L., and Hook-Barnard, I. (2025). Strengthening a Safe and Secure Nucleic Acid Synthesis Ecosystem: Outcomes of EBRC Stakeholder Engagement. Tech. rep., Engineering Biology Research Consortium

Millett, P., Alexanian, T., Brink, K. R., Carter, S. R., Diggans, J., Palmer, M. J., et al. (2023). Beyond biosecurity by taxonomic lists: Lessons, challenges, and opportunities. Health security 21, 521–529

of Sciences Engineering, N. A., Medicine, et al. (2018). Biodefense in the age of synthetic biology (National Academies Press)

Resnick, P. (2014). RFC 7282: On consensus and humming in the IETF. Online resource

SBOL Community (2025). SBOL community governance. Online resource https://sbolstandard.org/governance/

Tayouri, S., Kogan, V., Beal, J., Levy, T., Farbiash, D., Flyangolts, K., et al. (2025). Defending synthetic dna orders against splitting-based obfuscation. bioRxiv, 2025–03

The White House (2025). Executive Order on Improving the Safety and Security of Biological Research. Tech. rep., The White House. EO 14292

The White House Office of Science and Technology Policy (OSTP) (April 2024). Framework for nucleic acid synthesis screening. Online resource

UK Department for Science, Innovation, and Technology (2024). UK Screening Guidance on Synthetic Nucleic Acids for Users and Providers. Tech. rep., UK Department for Science, Innovation, and Technology

Warsaw, B., Hylton, J., Goodger, D., and Coghlan, N. (2000). PEP 1 – PEP purpose and guidelines. Online resource Python Enhancement Proposals, https://peps.python.org/pep-0001/

Wheeler, N. E., Bartling, C., Carter, S. R., Clore, A., Diggans, J., Flyangolts, K., et al. (2024a). Progress and prospects for a nucleic acid screening test set. Applied Biosafety

Wheeler, N. E., Carter, S. R., Alexanian, T., Isaac, C., Yassif, J., and Millet, P. (2024b). Developing a common global baseline for nucleic acid synthesis screening. Applied Biosafety 29, 71–78. doi:10.1089/apb.2023.0034

Wittmann, B. J., Alexanian, T., Bartling, C., Beal, J., Clore, A., Diggans, J., et al. (2025). Strengthening nucleic acid biosecurity screening against generative protein design tools. Science 390, 82–87

Wittmann, B. J., Wheeler, N. E., Murphey, S. T., Mitchell, T., Magalis, B., Gemler, B., et al. (2026). The limits of sequence-based biosecurity screening tools in the age of ai-assisted protein design. bioRxiv, 2026–03

World Health Organization (2022). Global Guidance Framework for the Responsible Use of the Life Sciences: Mitigating Biorisks and Governing Dual-Use Research. Tech. rep., World Health Organization

Wyschogrod, D., Manthey, J., Mitchell, T., Murphy, S., Clore, A., and Beal, J. (2022). Adapting malware detection to dna screening. In International Workshop on BioDesign Automation (IWBDA)

