## Supplementary File 1 for "Developing a Standard Definition for Sequences of Concern"

### RCP 001: Rubric Change Proposals

| RCP | 001 |
| --- | --- |
| **Title** | Rubric Change Proposals |
| **Authors** | Jacob Beal |
| **Type** | Process |
| **Status** | Active |
| **Discussion Issue** | #1 |
| **Rubric Changes** | PR #2 |
| **Test Set Changes** | N/A |
| **Created** | 29-Oct-2024 |
| **Approved** | 11-Dec-2024 |

#### Abstract

RCP stands for Rubric Change Proposal. An RCP is the means by which the Sequence Biosecurity Risk Consortium (SBRC) makes and documents decisions. The primary purpose of RCPs is to propose changes to the SBRC's Biosecurity Flag Rubric for assessing whether or not a biological sequence should be flagged as a potential sequence of concern. RCPs may also propose changes to the community's processes regarding RCPs.

The RCP should provide the rationale and a concise technical specification of the feature.

We intend RCPs to be the primary mechanisms for proposing such changes, for collecting community input on issues, and for documenting the design decisions that have gone into the BFR. As such, an RCP should provide the rationale for and a concise technical descriptiong of the changes proposed, along with examples of the consequences of the change and links to reference material and pending pull requests to implement the change. The RCP authors are responsible for building consensus within the community and documenting dissenting opinions.

RCPs are filed in the RCP directory of the central repository for the Screening Testing repository, and are accompaniede by an issue on the same repository for dicsussion of the RCP. When the authors of an RCP believe that rough consensus has been achieved by the stakeholder community, that consensus will be affirmed by a formal approval poll.

#### Table of Contents


- 1. Rationale
- 2. Specification
  - 2.1 RCP Workflow
    - 2.1.1 Drafting an RCP
    - 2.1.2 RCP Discussion and Decision-Making
    - 2.1.3 Application
    - 2.1.4 Minor Changes
  - 2.2 The RCP document
    - 2.2.1 RCP Types
    - 2.2.2 RCP Status
    - 2.2.3 Document Layout
    - 2.2.4 Auxiliary Files
- 3. Backwards Compatibility
- 4. Interaction With Other RCPs


#### 1. Rationale

A key challenge for the Sequence Biosecurity Risk Consortium (SBRC) is to be able make decisions about how to assess sequence risk both rapidly and also with sufficient stakeholder involvement to ensure that these risk assessments are widely acceptable. Since it is difficult to convene stakeholders for real-time discussions, it is important to be able to support asynchronous discussion and decisions-making. Likewise, it is important to have clear summaries of proposed decisions as well as readily accessible records of both the justification for a decision and the process by which it was made.

Rubric Change Proposals (RCPs) aim to address these issues. The RCP process is borrowing heavily from both Python Enhancement Proposals (PEPs), which are the main avenue to proposing and managing changes to the Python programming language, and SBOL Enhancement Proposals (SEPs), which are the main means by which the SBOL Data and SBOL Visual standards have been maintained. These processes have been used and refined by their respective communities over many years and we hope to benefit from this experience.

RCPs have the following goals:

- Manage proposed changes to the Biosecurity Flag Rubric (BFR).
- Enable lightweight changes to sequence test sets by application of the BFR.
- Distinguish between concrete proposals versus informal suggestions and aspirational discussions.
- Summarize arguments **for** and **against** a proposed change.
- Introduce newcomers and bystanders to an issue under discussion.
- Provide a framework for determining and affirming when approximate consensus has been achieved.
- Document history of decision-making in the SBRC community.

By drafting proposals as a shared document, the RCP process is intended to help integrate the diverse opinions and perspectives presented in a discussion thread. It encourages consensus-making by forcing co-authors of a proposal to make their change proposals concrete and grounded in examples, as well as to consolidate their opinions around the strongest points of agreement while resolving minor points of contention. Authors of a proposed change will be motivated to acknowledge, summarize, and address contrary points of view other than their own.

### 2. Specification

#### 2.1 RCP Workflow

##### 2.1.1 Drafting an RCP

The RCP process begins with a new idea for improving the BFR or its associated processes. It is highly recommended that a single RCP contain only a single key proposal or new idea. The more focused the RCP, the more successful it will tend to be, while broad and unfocused RCPs may have difficulty achieving approsimate consensus If in doubt, split your RCP into several well-focused ones.

The RCP author or authors write the RCP using the style and format described below, shepherd the discussions in the appropriate forums, and attempt to build community consensus around the idea. The authors are strongly encouraged to first socialize the idea with the community, either through dicsussion in meetings or on a GitHub issue, in order to determine if there is likely to be a viable path to consensus.

The authors then write up a plain text document, based on the template provided as RCP 002, summarizing their proposal as succinctly and clearly as possible (see below for details).

The authors should also draft a corresponding pull request containing the actual modifications that will be made to the BFR or any other documentation if the RCP is accepted. If the BFR is being changed, the authors should also create at least one pull request recategorizing sequences based on the proposed changes to the BFR. The purpose of these associated pull requests is to ensure that the proposal is fully defined and, in the case of rubric changes, to demonstrate that the proposed change is both necessary and practical.

Finally, when a draft is ready for discussion, the RCP should be submitted to the Screening-Test-Set repository, adding it to the `RCPs` directory, and associating it with a linked GitHub issue for discussion. If the RCP applies only to a specific cluster, then follow the same procedure, only in the repository for the cluster, rather than the `Screening-Test-Set` repository.

Note that merging an RCP into the main branch of the repository should happen at the start of the discussion, not wait for acceptance: RCPs provide a record of all serious proposals considered by the community, whether or not they are ultimately accepted.

###### 2.1.2 RCP Discussion and Decision-Making

Authors should update their RCP as the discussion progresses and their ideas are refined. Such updates should include documentation of alternative and dissenting opinions. Ideally, the authors should handle all such documentation themselves, but if the authors are not sufficiently responsive, other community members can propose their own amendments to the RCP discussion section.

Every RCP should eventually either be accepted or withdrawn. The decision to accept an RCP is made on the bases of rough consensus, following the IETF definition of there being no outstanding significant concerns that have not yet been addressed. As a practical matter, consensus can be achieved either in synchronous meetings or by quiescence of an asynchronous discussion thread.

Because many stakeholders may be infrequent participants who are not closely tracking community activity, however, rough consensus will be confirmed by means of an announced poll, whose purpose is to ensure that all relevant stakeholders have an opportunity to examine the proposal and raise any concerns they may have. As such, polling proceeds by the following process:

1. A poll is announced to the community, including a polling form with a link to the proposal and its associated discussion issue.

2. Announcement of the poll begins a 1 week comment period. During this time, if significant concerns are raised that cannot be readily addressed, then the poll should be cancelled.

3. After the comment period, the polling form opens for 1 week of feedback collection.

- Polling is confidential, but not anonymous: the people running the poll need to be able to see the identity of respondents who are objecting in order to be able to resume the search for consensus in the unlikely event that a poll fails.
- Poll responses can be "signed" by a person identifying themselves and their organizational affiliation.
- Poll options should be "Yes", "No", "Needs more study", and "Abstain"
- Polls should include an option for comments, so that people opposing adoption of a proposal can explain why they believe rough consensus has not been achieved.

If rough consensus is affirmed by poll results, the RCP will be marked as “Accepted”, and once its changes have been implemented, the status changes to “Final”. Approved procedural SEPs (e.g. concerning SBOL governing rules) are labelled as "Active", indicating that this rule may be further adapted in the future.

RCPs remain "Open" on the issue tracker until they no longer require attention; i.e., have been either accepted and merged, rejected, or replaced.

###### 2.1.3 Application

Upon acceptance, the pending pull requests linked in the RCP should be merged, thereby implementing the rubric change and its first application to recategorize test sequences.

In many cases, a rubric change will affect the categorization of other test sequences that are not contained in the pull requests for an RCP. For example, an RCP may affect many sequences, and only a representative subset are necessary for decision making, or some sequences may not be identified as affected by the RCP before it is implemented.

A key goal of the RCP process, however, is to make it fast and easy to recategorize test sequences, as long as the change is bringing their categories into compliance with the accepted rubric. As such, once an RCP has been accepted, it can be applied to recategorize test sequences by the following process:

1. Create a pull request that moves one or more sequences

2. In the pull request description, write down the application of the rubric that justifies the new categorization.

3. The pull request must be reviewed and approved, with the set of approvers containing at least:

    - 2 industry tool providers
    - 1 industry tool user

   - These organizations must be different from one another and from the organization of the pull request creator (i.e., at least 4 organizations involved in the pull request).

4. Pull requests must be open for at least 3 business days (balancing speed with time for people not working closely on the topic to have time to notice and review the request).

5. Any person can object to a pull request as being an incorrect application of the rubric, in which case, approximate consensus must be achieved or the rubric adjusted.

###### 2.1.4 Minor Changes

Some adjustments to the BFR or RCP processes are not significant enough to merit a full RCP process and a poll. For example, fixing typos, clarifying wording, or expanding documentation. In general, a change can be considered "minor" if the change is not expected to modify the classification of any test sequence or the substantive operation of any community process.

A minor change may be proposed and accepted using the follwing process:

1. Draft a pull request to modify the BFR or a process RCP.
2. Label the pull request as "Minor Change"
3. The pull request must be reviewed and approved following the same rules as for application of the RCP.
4. If any person objects to the labeling of the pull request as "Minor Change" and is not readily satisfied by discussion on the pull request, then the full RCP process must be used.

##### 2.2 The RCP document

###### 2.2.1 RCP Types

There are two types of RCPs:

- Rubric -- a proposal to change the Biosecurity Flag Rubric
- Process -- a proposal for improving processes related to RCPs

###### 2.2.2 RCP Status

Every RCP starts out in “Draft” status. A draft can become “Accepted”, “Rejected” or “Withdrawn”. A draft may also be “Deferred” if a discussion is deemed to be postponed.

After its implementation, an RCP is labelled as “Final”. Later during the life cycle of an RCP it may be “Replaced” or “Deprecated”. Process RCPs are instead labelled “Active”.

###### 2.2.3 Document Layout

An RCP is described as a text document using (GitHub-flavoured) Markdown syntax. A template for writing a new RCP is provided as RCP 002. This document, RCP 001, also follows the same template.

If necessary, authors may choose to change or introduce additional top-level sections but should follow this layout as closely as possible.

###### 2.2.4 Auxiliary Files

RCPs may include auxiliary files such as diagrams. Such files must be named `rcp-XXX-NAME.ext`, where `XXX` is the RCP number, `NAME` is the name of the auxiliary file, and `ext` is replaced by the actual file extension (e.g. `png`).

All auxiliary files should be linked from their RCP.

If there are more than 1-2 auxiliary files, they should be placed in a subdirectory named `rcp-XXX-files` to avoid cluttering the RCP list.

#### 3. Backwards Compatibility

As this is the first RCP, there are no backward compatibility issues to consider.

#### 4. Interaction With Other RCPs

- A template for proposing RCPs is given in RCP 002.
- A top-level rubric proposal to bootstrap the rubric is given in RCP 003
